# Fine-scale co-ocurrence patterns in grasslands reflect competition for space rather than broad plant strategies

**DOI:** 10.1101/2024.12.25.630317

**Authors:** Alexandre Génin, Louis Devresse, Eric Garnier, Sylvain Coq

## Abstract

1. Estimating the sign and strength of interactions among plants is central to understand the dynamics and functioning of communities, but is challenging to do for species-rich communities. Instead, spatial relationships between plants (clustering or spatial segregation) are sometimes used as a surrogate for the net effect of interactions ocurring between plants (positive or negative, respectively). However, this approach remains poorly tested outside of arid and alpine ecosystems, the ecological settings it originated from.
2. In experimental rangelands, we explored how management intensification, sheep exclusion and a natural soil depth gradient control the level of plant spatial segregation, or ‘negative co-occurrence’, usually considered as a measurement of competition intensity. We link these spatial patterns to classical broad plant strategies defined by 11 locally measured functional traits, and to the realized vegetation height and cover.
3. Plant segregation was highest when both grazing and fertilization were applied. Unexpectedly, general plant strategies (competitive, and acquisitive strategies) had little relationship with plant spatial patterns. Instead, spatial constraints increased segregation wherever cover was high and free bare ground was limited, or where plant growth is restricted by grazing to a few centimeters above ground.
4. These results show that fine-scale spatial patterns appear to capture competition for space, rather than for light or resources, as suggested by broad plant strategies. This may explain discrepancies in conclusions drawn from spatial patterns in grasslands, and clarifies the way towards a mechanistic understanding of spatial patterns.
5. *Synthesis*. The fine scale spatial organization of plant communities has been thought to reflect the intensity of competition among plants, but this approach has struggled to provide consistent results in grasslands. We show here that spatial patterns reflect competition for space rather than broad plant strategies captured by plant functional traits, helping us read observed plant spatial patterns to map interactions among plants in the field.

## Introduction

Identifying how plants interact with each other in nature, through positive or negative interactions, is central to our ability to determine the fate of ecological communities, and how they may respond to perturbations. While pairwise experiments have proven essential to quantify such interactions, both negative and positive, they are seldom implemented for species rich communities (Halty et al. 2017). The number of experiments to carry out becomes too high, and the interactions measured pairwise do not always reflect the effects species have on each other in the context of a complex ecological community (Wootton 1994, Engel and Weltzin 2008, Mayfield and Stouffer 2017, Levine et al. 2017).

A recent approach based on the spatial distribution of plant individuals has been proposed to estimate plant-plant interaction. Competing species are expected to inhibit the development of each other, and thus on average, they should be found segregated from each other in space. Conversely, plants facilitating each other are expected to establish preferentially close to each other, and thus be spatially aggregated. This approach has been used in arid ecosystems, where facilitative interactions lead to an unambiguous pattern of aggregation between nurse shrubs and facilitated species under their canopy (Verdú and Valiente-Banuet 2008). It has been extended to include both spatial aggregation and segregation, and thus both potential positive and negative interactions, with salient applications in abiotically-stressed ecosystems (Choler et al. 2001, Saiz and Alados 2011, Losapio et al. 2021). However, these studies rely on the fact that in those ecosystems, spatial patterns are distinct enough to be unambiguously attributed to species interactions, often positive, and experimental work has been carried out to show that this is effectively the case (e.g. Valiente-Banuet and Ezcurra 1991, Graff and Aguiar 2011). This is not true for many other ecological settings. In the case of negative interactions in grasslands, the link between spatial patterns and plant-plant interactions, when tested, is often tenuous or inconsistent with experimental results (Schamp et al. 2022). For example Brazeau and Schamp (2019) found that spatial segregation appears unrelated to negative effects of better competitors on weaker ones. Conversely, Gridzak et al. (2024) do observe that pairs of plants with large differences in height tend to segregate in space, pointing to a negative effect of taller plants. These discrepancies may arise because spatial patterns can be substantially modified by environmental disturbances, such as trampling, independently of plant-plant interactions (Saiz and Alados 2012, Alados et al. 2017, Génin et al. 2021). This comes on top of statistical limitations of the methods used to infer potential interactions from spatial patterns, which can lack statistical power (Rajala et al. 2019), or be sensitive to spatial scale or other confounding factors (McNickle et al. 2018, Blanchet et al. 2020).

We argue that a more mechanistic understanding of the ecological processes captured by spatial patterns is key to reconciling discrepancies between studies and refining the links between spatial patterns and plant-plant interactions. To do so, plant spatial patterns need to be related to the local characteristics of the vegetation that are potentially involved in interactions.

This includes the general vegetation morphology, which includes the total local vegetation cover and canopy structure. Competition for space, nutrients or light should be stronger in denser communities with a lower proportion of bare ground. The outcome and intensity of competition for space, is also largely controlled by plant growth forms, a functional categorisation based on the direction and extent of growth (Pérez-Harguindeguy et al. 2016). In grasslands, due to the high density of vegetative material near the soil surface, it is likely that rosette or tussock plants are on average better competitors for horizontal space than that of extensive and stemmed herbs.

Second, plant strategies captured by functional traits are also expected to be related to plant-plant interactions (Mcgill et al. 2006). Syndromes of traits are thought to capture to some extent the ecological strategy of plants, particularly in terms of resource acquisition. For example, species with higher competitive ability for light are expected to have a high maximum height. In addition, plants with low leaf dry matter content (LDMC) and high specific leaf area (SLA) acquire resources at a high rate and have a high relative growth rate. In certain contexts, such resource-abundant plants may be competitive. These two axes of variation, height and resource acquisition strategy, are well documented trait syndromes that are used to estimate the dominant type of strategy in a community in situ (Garnier et al. 2015, Díaz et al. 2016). Communities where competition for light is stronger are expected to have a greater proportion of taller plants in close proximity. Similarly, communities with high nutrient availability will generally have higher proportions of resource-acquiring strategies. Plant functional traits are therefore expected to be related, at least to some extent, to the strength of competition in the field.

The vegetation spatial patterns, morphology and functional composition all depend on environmental conditions and resource availability, and are therefore expected to be affected by land use. In rangelands, a gradient of intensification can be considered, involving different levels of fertilization and grazing intensity (Blüthgen et al. 2012). High nutrient availability is typically associated with tall and acquisitive plants, and allows for high aboveground biomass. High levels of competitive exclusion are expected to occur in this context, leading to spatially-segregated communities. By contrast, scarce soil resources favour plant communities with lower total cover, in which plants are short and have conservative strategies, as shown previously at our study site (Bernard-Verdier et al. 2012, Pérez-Ramos et al. 2012). The lower competition levels, or even the possible facilitation in those communities are expected to favour spatial aggregation between plants.

Grazing on the other hand is expected to have a more complex effect. First, grazing compounds with resource availability to shape the dominant strategy of plants in communities. Plants species persisting under high grazing intensity in relatively fertile areas are usually prostrate (rosette plants in particular are abundant), small, and have leaves with high SLA and low LDMC (Garnier et al. 2015). Biomass removal may also lead to a relaxation of competitive effects between plants, whereby the removal of tall species alleviates competition for light and allows subordinate species, which are shorter or not as resource-acquisitive, to persist in the community (Eskelinen et al. 2022). In addition to this effect on the plant community, herbivores directly shape the morphology of individual plants through trampling and removal of biomass. This may directly affect spatial patterns, and thus estimates of competition from those (Alados et al. 2017), as well as conclusions drawn from trait measurements.

To relate plant spatial patterns to relevant community characteristics, we used a long-term experiment with two levels of land use intensification (involving both fertilisation and grazing intensity), 18 fenced sheep exclosures, superimposed on a natural gradient of soil characteristics. We investigated how spatial patterns vary under these different conditions, and how they relate to field-measured vegetation characteristics, in particular to plant strategies captured by traits measured at the same site.

We hypothesized that:

- H1: Overall levels of competition, and thus spatial segregation, are higher in fertilized and ungrazed areas
- H2: Grazing decreases competition by reducing plant biomass, and thereby diminishes spatial segregation. This reduction is expected to be more pronounced in intensively-managed areas, where biomass production and removal are higher.
- H3: All else being equal, plant communities with fast acquisition strategies and taller plants should have show higher spatial segregation
- H4: Plant communities with higher amounts of growth forms that preempt horizontal space, such as tussocks or rosettes, should also show higher spatial segregation. These effects should be particularly strong where available space is low, i.e. where vegetation cover is high or plants have very similar heights.

## Methods

### Environmental gradients at La Fage

Our study site is located at the *La Fage* Experimental Station (43°55′N, 3°06′E, 765–830 m a.s.l.), a facility of the French National Institute for Agricultural and Environmental Research (INRAE), in an area traditionally subjected to extensive sheep grazing. This long-term experimental study site for grazing systems has hosted a number of studies on the effects of grazing and intensification of agricultural practices on the functional structure of plant communities (Bernard-Verdier et al. 2012, Pérez-Ramos et al. 2012, Garnier et al. 2018, Coq et al. 2018). The main experimental factors at La Fage include a fertilization treatment: since 1978, some paddocks have been fertilized with inorganic nitrogen (65 kg NH_4_NO_3_.ha^-1^ every year) and phosphorus (40 kg H_3_PO_4_ .ha^-1^ every three years until 2003), while others have been left unfertilized. Grazing pressure is also higher in fertilized areas, with a higher proportion of the total annual biomass produced consumed by sheep (0.61 kg.kg^-1^) compared to native areas (0.20 kg.kg^-1^). In March 2012, sheep exclosures were established in three fertilized and three unfertilised grazed paddocks. Three 10×10 m exclosures were set up in each paddock, for a total of 18 exclosures (Coq et al. 2018).

These experimental factors are overlaid on a natural gradient of soil depth (Bernard-Verdier et al. 2012, Pérez-Ramos et al. 2012). Some areas of the study site have shallow soils where coarser material from the underlying bedrock makes up a significant proportion of the soil, while others have deeper soils made up of finer material (doline-like areas with silt). We captured the effect of this natural soil gradient by visually estimating the cover of bare ground, coarse pebbles (> 5 mm), fine pebbles (< 5 mm) and apparent bedrock for each sampled site in ten 50×50 cm quadrats regularly spaced on either side of a 2.50 m linear transect. These covers were then used as input for a principal component analysis and we extracted the first axis to obtain a “soil depth index”, which explained 77 % of the variance (Supplementary Figure S1).

### Defining plant strategies from traits

Nine traits were selected to capture plant strategies (Figure S2; Table S1), using data from previous studies carried out at our study site (Table S1; Bernard-Verdier et al. 2012, Pérez-Ramos et al. 2012, Chollet et al. 2014). Where available, trait values were obtained from the fertilized and unfertilized treatments on individuals not affected by grazing. For 90 % of species and trait combinations, the trait average was made with a least 4 values. Trait syndromes were constructed from the data by taking the average trait measured for each species and using it as input for a Principal Component Analysis (PCA). This allowed us to identify two main axes of variation previously identified at our study site (Pérez-Ramos et al. 2012, Garnier et al. 2018), reflecting the slow-fast continuum of leaf economy (Axis 1) and the ability to compete for light (Axis 2), which explained in total 61 % of the trait variability in the dataset (Figure S2).

### Characterizing levels of spatial segregation

We characterized the spatial structure of plant communities based on 2.50 m long transects. For each transect, we laid out a tape line and recorded the start and end of the length over which any plant vegetative part (i.e. leaf or stem) overlaped the line, to an accuracy of 2 mm (Figure S3; Génin et al. 2021). Transects were carried out at 18 different locations in the study site. At each location, we carried out four transects to measure species associations, composition and soil depth: two in an exclosure (ungrazed treatment) and two a few meters from the exclosure.

From this dataset, for a given transect, we calculated the total observed overlap between each species Oij (Figure S3) This overlap was then compared to a null distribution obtained by randomizing 2999 times the position of each individual segment along the transect. This process yields a set of association strengths between all species occurring within a transect. We considered a negative association to occur between two species if the observed overlap O_ij_ was less than 75% of the null distribution (i.e. *α* =0.25). We then summed all negative associations to obtain the raw total number of negative associations K. This protocol is the direct equivalent for continuous sampling of defining associations based on the number of sampling sites shared between two species (Saiz and Alados 2012, Sanderson and Pimm 2015).

Along environmental gradients, species abundances change, which affects our ability to identify negative associations (Saiz and Alados 2012). For example, it is impossible to detect that two species with very low cover are spatially segregated because their observed overlap Oij will be close to or exactly zero, which is also what is expected by chance. As the number of such low-cover species varies between sites, the total number of negative associations K will vary, even in the absence of changes in spatial patterns. Therefore, it is necessary to correct for this effect and remove the confounding effect of changing covers when comparing spatial associations between different sites (transects). We did so by quantifying how much K deviates to what is expected given the distribution of species covers in the plant community. To do this, for a given transect, we shuffled the positions of the plant individuals along the transect 999 times and recalculated the number of negative associations as above, this time starting from this randomized data set. For a given transect, this resulted in a null distribution for the number of negative associations K, which we summarized into a mean *μ_null_* and standard deviation *σ_null_*. The corrected number of negative associations was then calculated as *K_SES_*=(*K* −*μ_null_*)/ *σ_null_*.

KSES quantifies by how much the observed number of negative associations is above or below its null expectation, and can be compared across transects with different species covers and richness. It is, by construction, independent of these two variables (whereas K is not). A positive value means that there is an excess of negative associations, and a negative number a deficit in negative associations. A value of zero means that the observed number of negative associations is consistent with random expectation, i.e. what would be obtained given a random distribution of plants in space. We obtained one value for K_SES_ per transect, which resulted in 72 values in total (6 per treatment).

### Characterizing the dominant strategies per treatment

To investigate how the functional structure of communities varied with natural gradients and experimental treatments, we computed for each transect the community-weighted mean (CWM) for each trait. This provides a single number that reflects the average trait value in a given plant community, weighted by the total abundance of each species. We used trait measurements specifically measured in fertilized or unfertilized areas, in other words CWMs computed for fertilized sites were done so only based on trait measurement done in fertilized areas (and viceversa for unfertilized sites). 80 % of the transects had a trait coverage above 60 % (i.e. 60 % of the total species cover in transects had matching trait values).

The CWMs of a transect represents the “average species” found in this community in terms of functional traits. We used these values to infer the position of this average (virtual) species in the PCA done previously at the species level (Figure S2), to obtain two indices reflecting the typical species found in each transect in terms of fast-acquisition strategies (increasing values of Axis 1), or ability to compete for light (increasing values of Axis 2).

To visualize how the functional composition varied with the experimental treatments, we used a principal component analysis on the matrix of CWMs (transects as rows and traits as columns). To assess whether individual treatments had a significant effect on functional composition, we used a redundancy analysis (RDA) with fertilization, grazed status and soil fertility as predictors. We tested the significance of these predictors based on permutation tests (implemented in the R package “vegan”; Oksanen et al. 2018).

Finally, we measured the realized morphology of vegetation in the field based on field-recorded plant height. We estimated visually the height of each recorded plant using a categorical scale from 1 to 5, based on the following height intervals, in cm: [0, 2), [2, 5), [5, 15), [15, 30), [30, 60). We then computed for each transect the average height using class midpoints, weighted by the cover of each individual. We computed the percentage of transect length covered by plants in each transect, as a variable characterizing the amount of horizontal space used by plants (and thus the lack of space available for colonization). We used simple t-tests to assess the effect of grazing on the cover and average height of vegetation.

### Effects of environment, functional composition and plant morphology on negative associations

We used bayesian linear mixed-effects models to tease out the direct effects of Grazing, Fertilization, Soil depth, dominance of competition for light and fast acquisitive strategies in the plant community. We first used the number of negative associations as a response (KSES), and all of the above variables as predictors. We considered all combinations of interactions between categorical variables (Fertilization and Grazing) and continuous variables (Competition for light and Fast growth and Soil depth), and used a leave-one-out cross-validation criterion to discard interaction terms until the best balance between model predictive ability and overfitting was achieved (Vehtari et al. 2017). The best model according to this criterion only included interactions between the grazing status and fertilization.

We used a similar strategy based on model selection to estimate the explanatory power of realized vegetation morphology (height) and total plant cover on spatial patterns. We used *K_SES_* as a response, with the levels of grazing and fertilization (with interactions), soil fertility, average transect height and percentage cover. We dropped first all interaction terms as done above. However, we went further in terms removal to assess whether spatial patterns could be well-predicted only by morphological attributes of the vegetation (average height and cover), by considering models that excluded treatments as explanatory variables, on top of interactions. Again, we used leave-one-out cross validation to select the most parsimonious model out of these combinations, which was one that included only plant height and percent cover as explanatory variables.

In all bayesian linear models, we used the location as a random effect on the intercept, to take into account the repeated sampling in nearby areas. Analyses were performed using the packages vegan v2.6.4 (Oksanen et al. 2018), brms v2.21.0 (Bürkner 2018) in R (v4.4.1; R Core Team 2024). Throughout this manuscript, we use significance levels of alpha = 0.1 when using frequentist statistics, and 90% credible intervals when working with bayesian statistics, to take into account the reduced number of unique replicates per treatment combination (N = 3). We used uninformative, wide priors carrying out the bayesian regressions, namely N(0, 5) for explanatory variables, N(0, 10) for intercepts, and exp(5) for coefficients characterizing random effects or residual error.

## Results

### Environmental gradients

Based on the PCA done on the CWM of traits in each transect, we found that the functional composition of plant communities was mostly driven by the fertilization status (Figure 1a), while grazing appeared to have a smaller effect (Figure 1b) when considering sites altogether. The gradient of soil depth appeared to explain some of the variability in functional composition, and was very variable in unfertilized sites, but less so in fertilized sites (Figure 1c). These visual patterns were confirmed by permutation-based tests, which supported a significant effect of fertilization and soil depth index (RDA-based permutation tests, P<0.001), but only a marginal effect of grazing (P=0.10).

**Figure 1.**
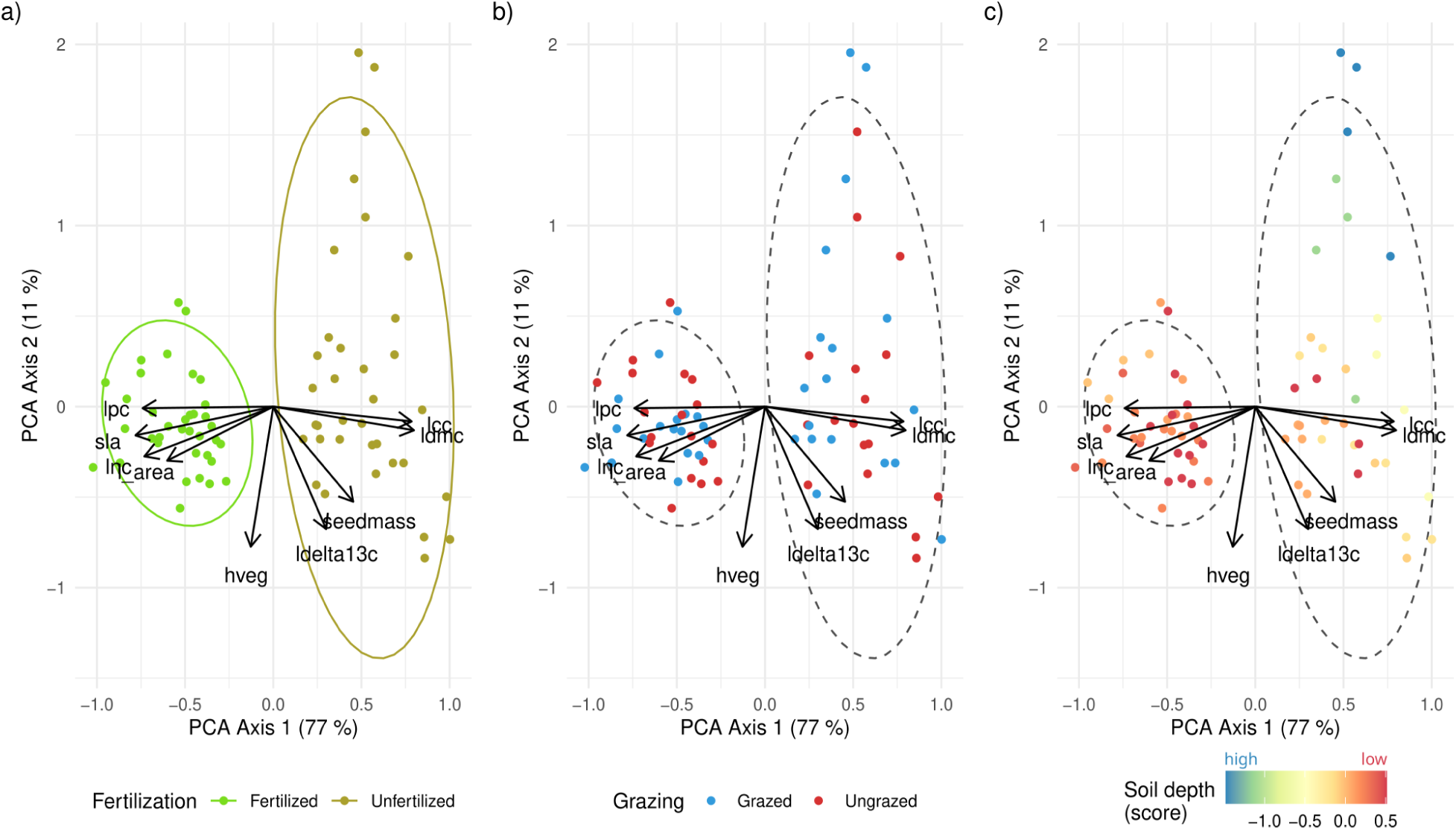
Effect of experimental treatments on the functional composition of plant communities. Each dot indicates a transect, whose coordinate in PCA space reflects its functional composition, as quantified by CWMs. Colors indicate in panel a), the fertilization status, in b) the grazing status, and in c) the soil depth score (higher score indicate higher soil depth). Ellipses indicate the 90% normal ellipses for fertilized and unfertilized points. See Table S1 for a complete description of functional traits and their abbreviations.

### Drivers of species associations

Linear modeling (Figure 2) revealed that soil depth and the dominant trait syndromes found in a transect (in terms of resource acquisition strategy and competitiveness) appeared to have no strong effect on the number of observed negative associations. K_SES_ was not explained by the average amount of acquisitive species in a transect, with a 90% credible interval for the corresponding model coefficient largely overlapping zero ([-1.07, 1.15]; see also full posterior distributions in Figure S3). Similarly, a larger amount of competitor species in a community did not appear to correlate with an increased number of negative associations (90% interval of [-0.66, 0.23]).

**Figure 2.**
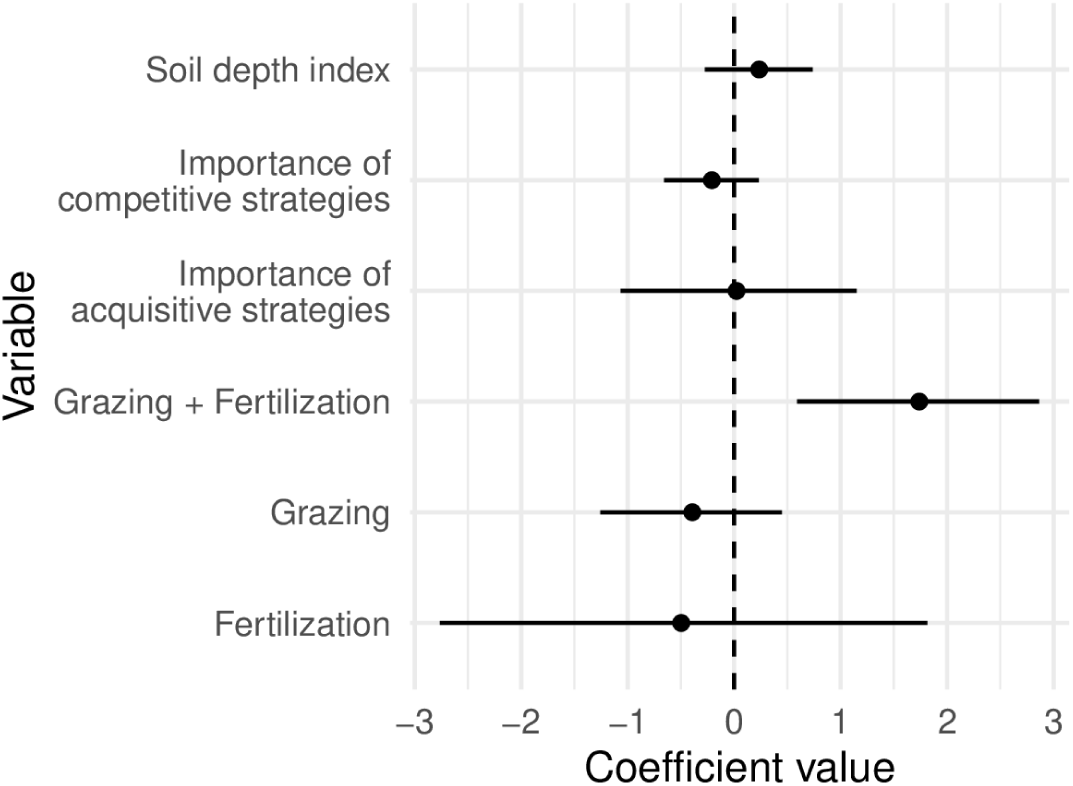
Estimated average and 90% credible intervals for the linear model including experimental treatments and importance of plant strategies in the transect as explanatory variables, and K_SES_ as response variable.

Grazing and fertilization had a discernible effect when applied together, with a 90% credible interval that did not include zero for this combination ([0.59, 2.87]). However, no equivalent effect of those treatments in isolation was found, as credible intervals largely overlapped zero ([-2.76; 1.82] and [-1.26; 0.45] for Grazing alone and Fertilization alone). This pattern was also suggested by post-hoc analyses (Figure 3), with only the Fertilized+Grazed treatment being significantly different from the other treatments (pairwise t-test with Benjamini-Hochberg correction, P < 0.1).

**Figure 3.**
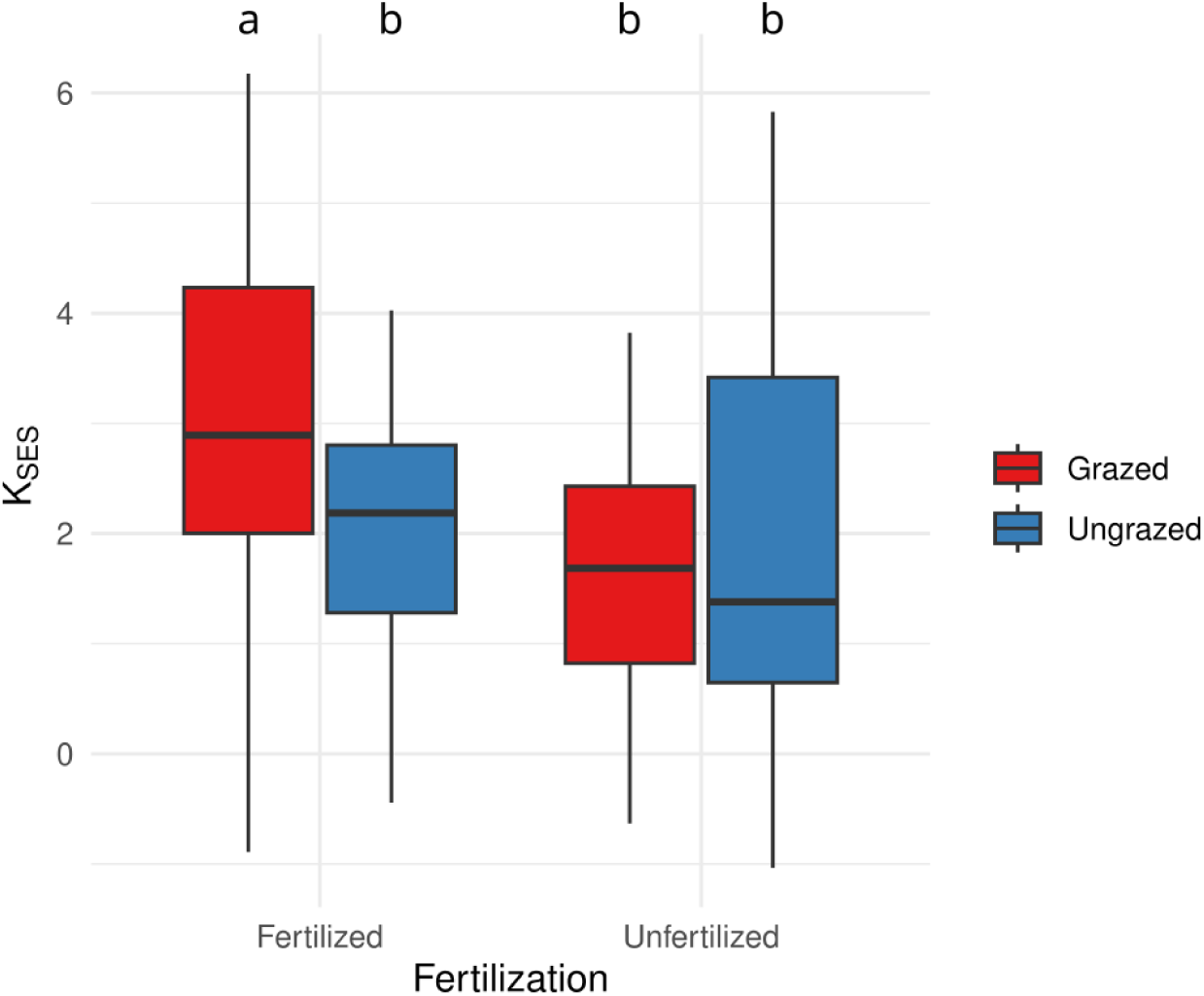
Negative associations observed for the different grazing and fertilization levels. The letters indicate the different groups, based on significant differences resulting from a pairwise t-test with Benjamini-Hochberg correction for multiple testing (P<0.1).

### Effects of plant morphology and available space on spatial structure

Turning now to the realized morphology of plant communities in the field, we found that grazing significantly reduced vegetation height (t-test yielding P<0.01; Figure 4), both in fertilized areas (3.0 ± 1.4 cm vs. 13.2 ± 6.7 cm; mean ± s.d.) and unfertilized areas (4.6 ± 3.5 cm vs. 10.7 ± 5.0 cm). However, grazing did not significantly reduce the percentage of soil covered by plants (P>0.1; Figure 5), both in fertilized (72 ± 13 % and 67 ± 9 % for grazed and ungrazed conditions, respectively), and unfertilized areas (51 ± 13 % and 54 ± 8 %).

**Figure 4.**
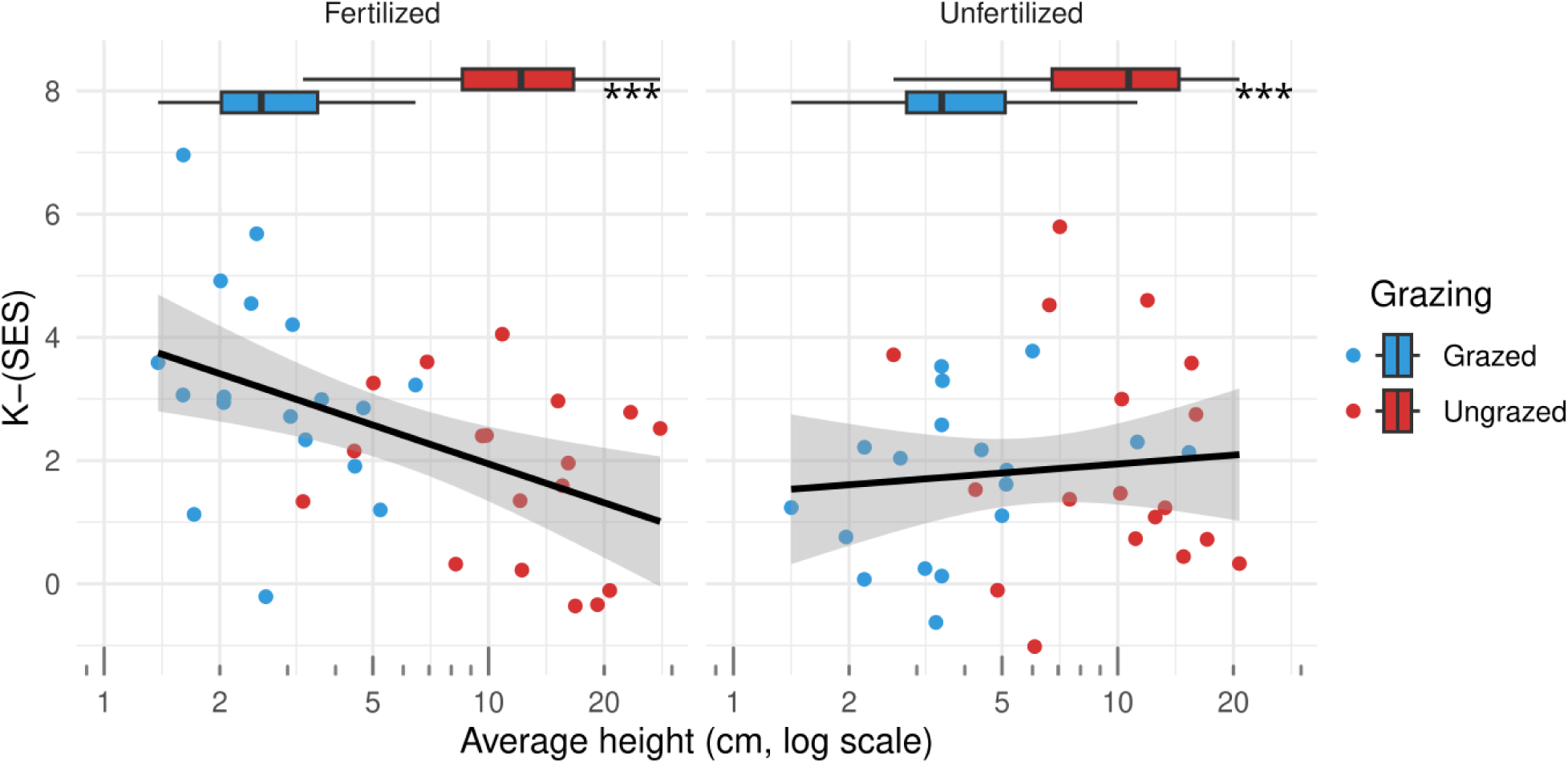
K_SES_ as a function of the average height (log scale), for fertilized (left panel) and unfertilized areas (right panel), and different grazing status (point color). The trend is a linear trend computed using all points in the relevant panel. Boxplots in the margin indicate the distribution of heights for each grazing treatments, with three asterisks indicating significant differences according to a t.test (P<0.01).

**Figure 5.**
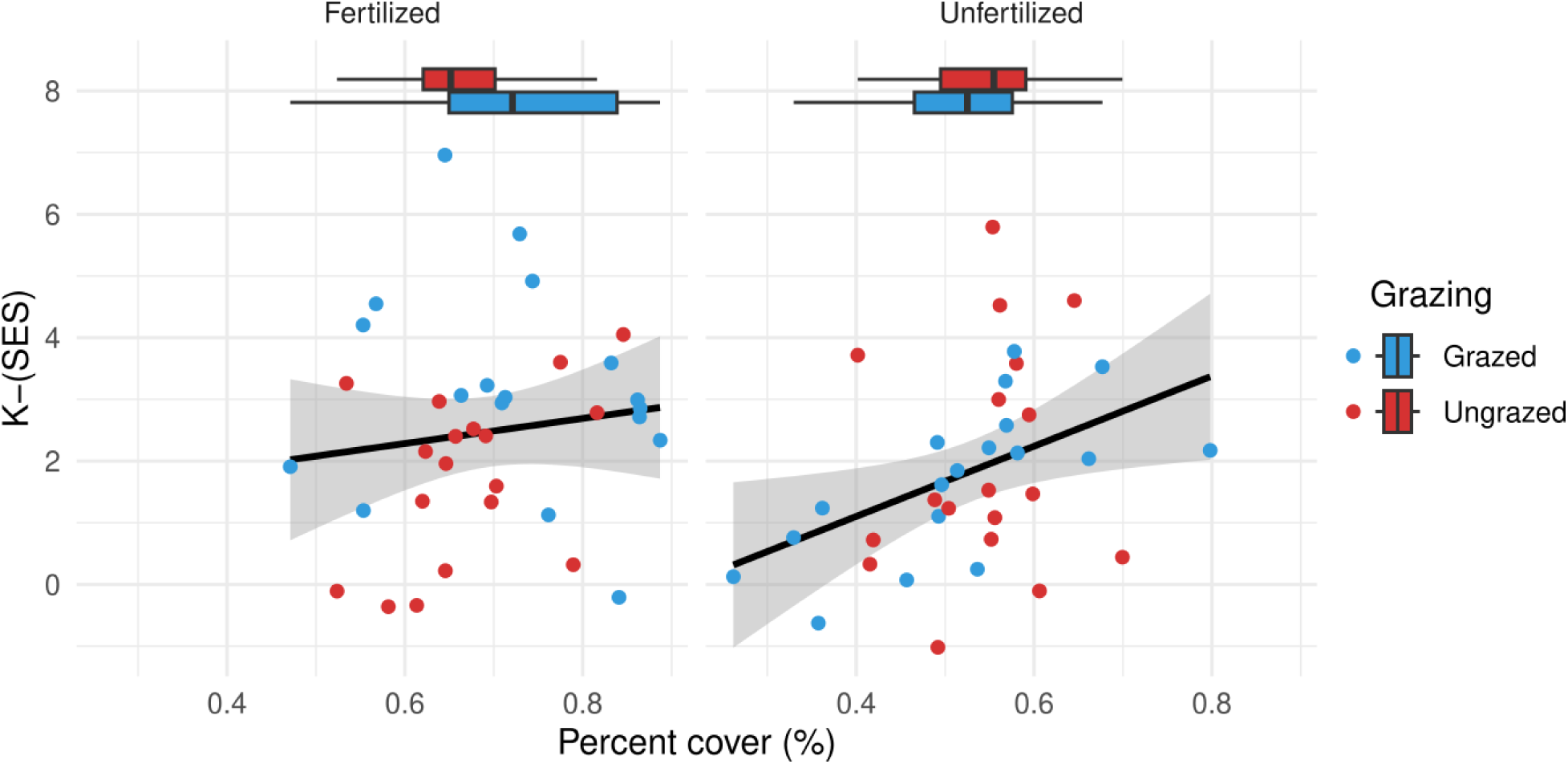
K_SES_ as a function of the total cover of plants in the transect, for fertilized (left panel) and unfertilized areas (right panel), and different grazing status (point color). The trend is a linear trend computed using all points in the panel. Boxplots in the margin indicate the distribution of covers for each grazing treatments (differences according to a t.test were non-significant, P>0.10, so no asterisks are displayed).

A linear model suggested that K_SES_ could be well-predicted only with total cover and average plant height. Both terms had an effect on K_SES_, as the model suggested a decrease of K_SES_ with average community height (90% credible interval of [-0.66, -0.09]; Figure 4), and increased with the total cover of plants in the transect ([0.22, 0.81]; Figure 5). Full posterior distributions for each coefficients are given in Figure S4.

## Discussion

As found in other studies focused on fine-scale spatial patterns in grasslands, our results confirm that large variations exist in the levels of plant segregation (K_SES_). In our case, these variations appeared linked to changes in environmental factors, though their effects was synergistic rather than additive, as grazing and fertilization had significant effects, but only when both were applied. This contradicts previous findings (Génin et al. 2021) and our initial hypotheses of significant effects of grazing and fertilization in isolation (H1, H2). Broad axes of variation in plant strategies defined on non-damaged individuals generally explained little variation in spatial patterns, again contradicting our initial expectation (H3). The realized morphology of vegetation however, both horizontally in terms of transect length covered by plants, as well as vertically, as measured by average plant height, were significant predictors of spatial patterns (H4).

Taken altogether, these patterns suggest that environmental factors do not control the spatial segregation of plants through direct effects, but by their effects on the morphology of vegetation. Spatial segregation was only significant under fertilized and grazed conditions, which coincided to areas where the development of plants was restricted to a few centimeters aboveground. This limits the ability of plants to overlap with each other, as they can only coexist side-by-side in the space available to them. Thus, grazing most likely increased spatial segregation by constraining plants so that they could only develop laterally. It is interesting to note that grazing did not give rise to such effect in unfertilized areas, possibly because vegetation there remained slightly taller despite grazing, compared to fertilized areas (typically 4-5 cm vs. 3 cm on average). This may be also explained by differences in total cover, as communities with little horizontal space left for colonization also had larger levels of spatial segregation. This occurred primarily in fertilized areas with higher productivity, where horizontal cover reached its highest values observed (around 90 %), and was overall higher by 20 % than in unfertilized areas. Grazed and fertilized areas had thus higher spatial segregation most likely due to additive effects of both vertical and horizontal constraints limiting the possible overlap of plants with each other.

Broad plant strategies defined on undamaged individuals were overall weak predictors of spatial segregation. This highlights that potentially competitive or acquisitive plants may not always show spatial patterns in relation to their expected trait syndrome, warning against interpreting spatial patterns in the light of expectations based on the common two-axes view of plant strategies. It is important to note that this does not mean that there is no relationship to expect between traits and spatial patterns, but this may rather concern traits not directly tied to height or leaf economy. For example, the type of life form, such as whether plants produce rosette phenotypes with leaves close to the ground appear to be a predictor of segregation, particularly in fertilized areas (Figure S6). Similarly, belowground traits may be more relevant to spatial patterns. Rhizomes and roots, as well as their symbionts, can contribute to the spatial organization of plant communities, and aerial parts of plants do not necessarily reflect that of belowground parts (Kesanakurti et al. 2011). Belowground spatial patterns are complex and change along soil layers. Consequently, the observed patterns may vary with soil conditions at our study site. In a pot study for example, Semchenko et al. (2018) demonstrated that different belowground traits relate to the ability of different species to tolerate competition and to dominate their competitors. *In-situ* measurements of belowground traits could be better linked to spatial patterns in our site. This option is however hardly feasible for the vast number of individuals involved in the evaluation of spatial patterns.

Beyond the type of trait measured, and despite the quality of our database, it is important to note that our ability to measure the effect of grazing on the functional composition of the community is limited here because traits were only measured on healthy individuals outside of exclosures. We measured traits for the same species in fertilized and unfertilized areas, so we could take into account any intraspecific change in plant strategy due to fertilization. We could detect for instance that a species had a larger potential vegetative height in fertilized areas, and thus becomes a better competitor for light as a result of fertilization, and carry this intraspecific change into the computation of CWMs. However, because we did not have similar measurements of traits inside exclosures, all functional changes between grazed and ungrazed treatments can only arise from species turnover. While species turnover did occur with exclosures, with typically half of the species replaced inside vs. outside the exclosure (Figure S7), this limits our ability to detect changes in plant strategies due to grazing, relative to changes in fertilization. Despite this limitation, it is unlikely that a large effect of plant strategies would have remained undetected, but further studies may be able to detect such subtle effect.

The spatial patterns measured here mostly reflect competition for space between plants – wherever space was restricted, either vertically or horizontally, our metric of spatial segregation increased. This partly arose because our sampling design considered two species as co-ocurring only when they were growing on top of, or below each other. This definition necessarily limits the number of individuals that can co-occur at a given location, because plants need specific combinations of morphologies to allow vertical overlap, for example because one of the pair is taller and allows smaller plants in its understory, or because one of them is a vine that can grow on other plants. Such effect is not present when there is no such physical constraint to plants growing in the same sampling unit, for example because sampling units are much larger than the size of an individual, or because plants are growing in very different vegetation strata (e.g. tall bushes vs. seedlings growing underneath). Interestingly, in such cases, spatial aggregation or segregation could better reflect other components of plant-plant interactions, such as competition for resources, or be less related to interactions altogether, and more to environmental drivers (Bektaş et al. 2023). The fact that spatial patterns may reflect different ecological processes depending on the sampling scale (McNickle et al. 2018) might explain why analyses based on co-occurrence yield inconsistent results in grasslands.

Using spatial patterns to investigate plant-plant interactions has recently seen a rise in interest as it allows mapping complex spatial association networks between plants, which are made of positive or negative links between species that represent spatial aggregation or segregation (Losapio et al. 2019). In this study, although we do not make use of the topology of such networks, our index of spatial segregation is in fact based on a tally of negative links. Plant-plant networks are rich descriptions of communities, as their structural properties, such as connectance or structural balance, can be investigated along time or environmental gradients to better understand the response of plant communities to disturbance (Saiz et al. 2018, Génin et al. 2021, Losapio et al. 2021). However, going beyond such descriptive analyses requires understanding how these networks of spatial relationships relate to interactions between plants in the field. Our study show that fine-scale spatial patterns appear to capture particularly well competition for vertical and horizontal space, but this may be different depending on the chosen methodology and species involved. These discrepancies hamper the efforts to develop a unifying theory of the structure of complex plant communities across ecosystems, as is done for example with food webs, which enable comparative analyses of ecological communities with very different species (Brose et al. 2006), and whose structure is governed by a narrow set of traits (e.g. body size; Williams and Martinez 2000). Reaching similar levels of unification for plant networks will require a much-improved understanding the nature of plant-plant interactions captured by spatial patterns.

## Acknowledgements

This work has benefited from invaluable feedback, financial support, and insights from Johanne Nahmani, Sonia Kéfi and Alain Danet. It is based on numerous years of field work carried out at INRAE Fage experimental station, whose staff we warmly thank.

## Supplementary material

**Figure S1.**
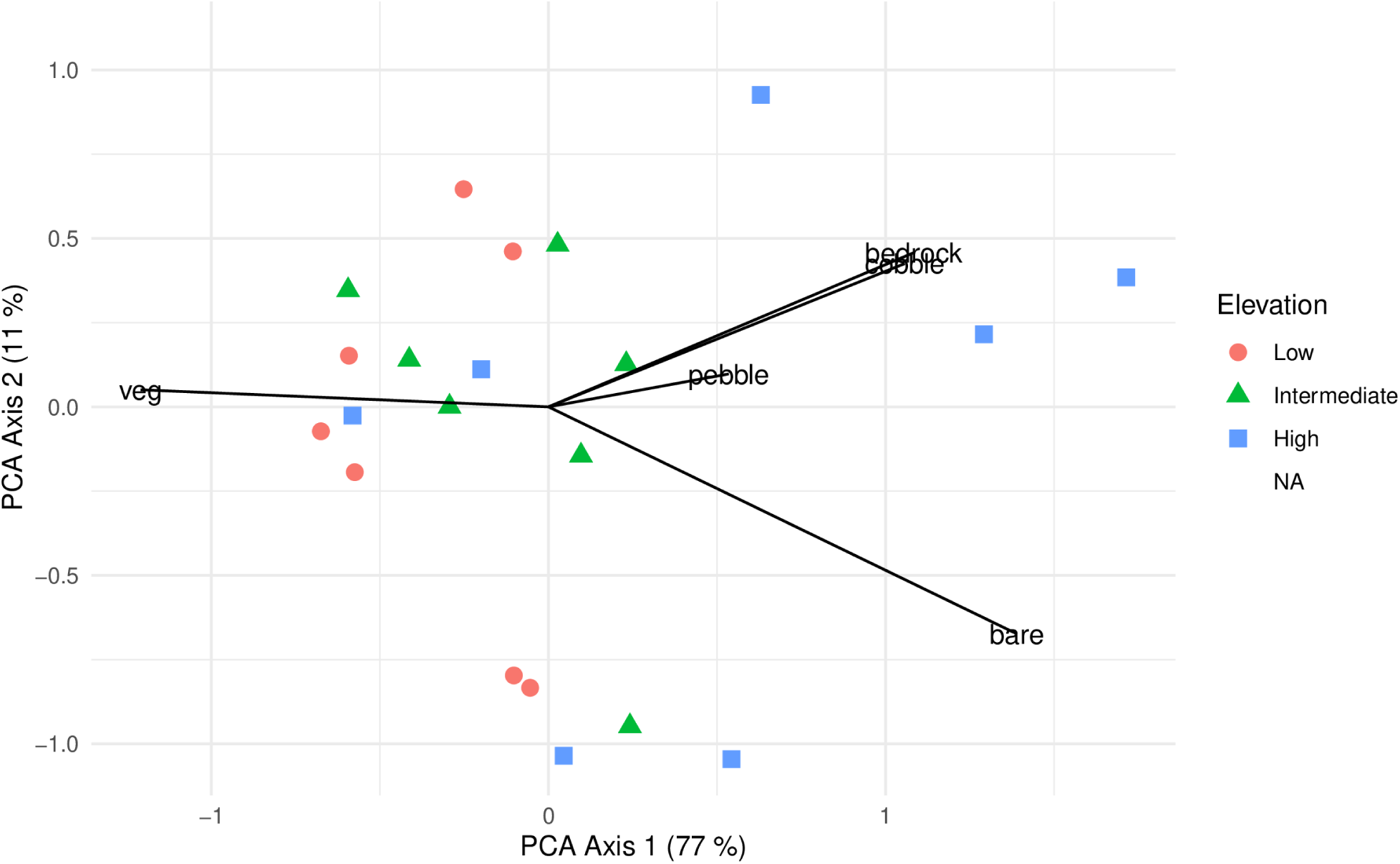
Principal Component Analysis on the 5 soil surface characteristics used to define the Soil Fertility Index. Arrow labels stand for % vegetation cover (veg), % exposed bare soil (bare), % pebble (areas covered with particles between 2 mm and 2 cm), % cobble (particles between 2 cm and 10 cm) and % bedrock (partly exposed rocks whose size is too big to be determined).

**Figure S2.**
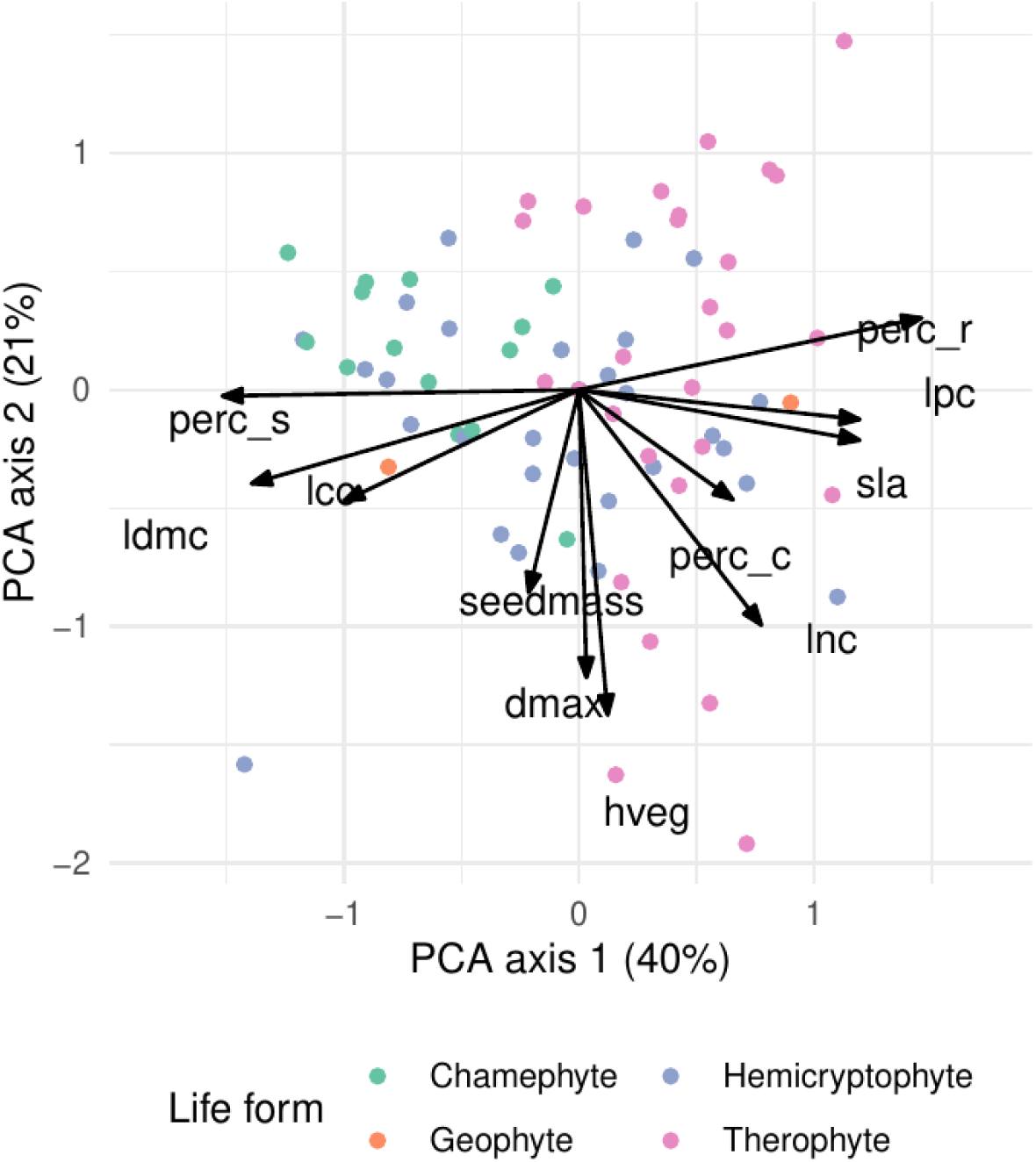
Principal Component Analysis (PCA) results on species. In this graph, each point represents a species, and its average trait value is used as input to the PCA. Point colors indicate the Raunkier life form for each species.

**Figure S3.**
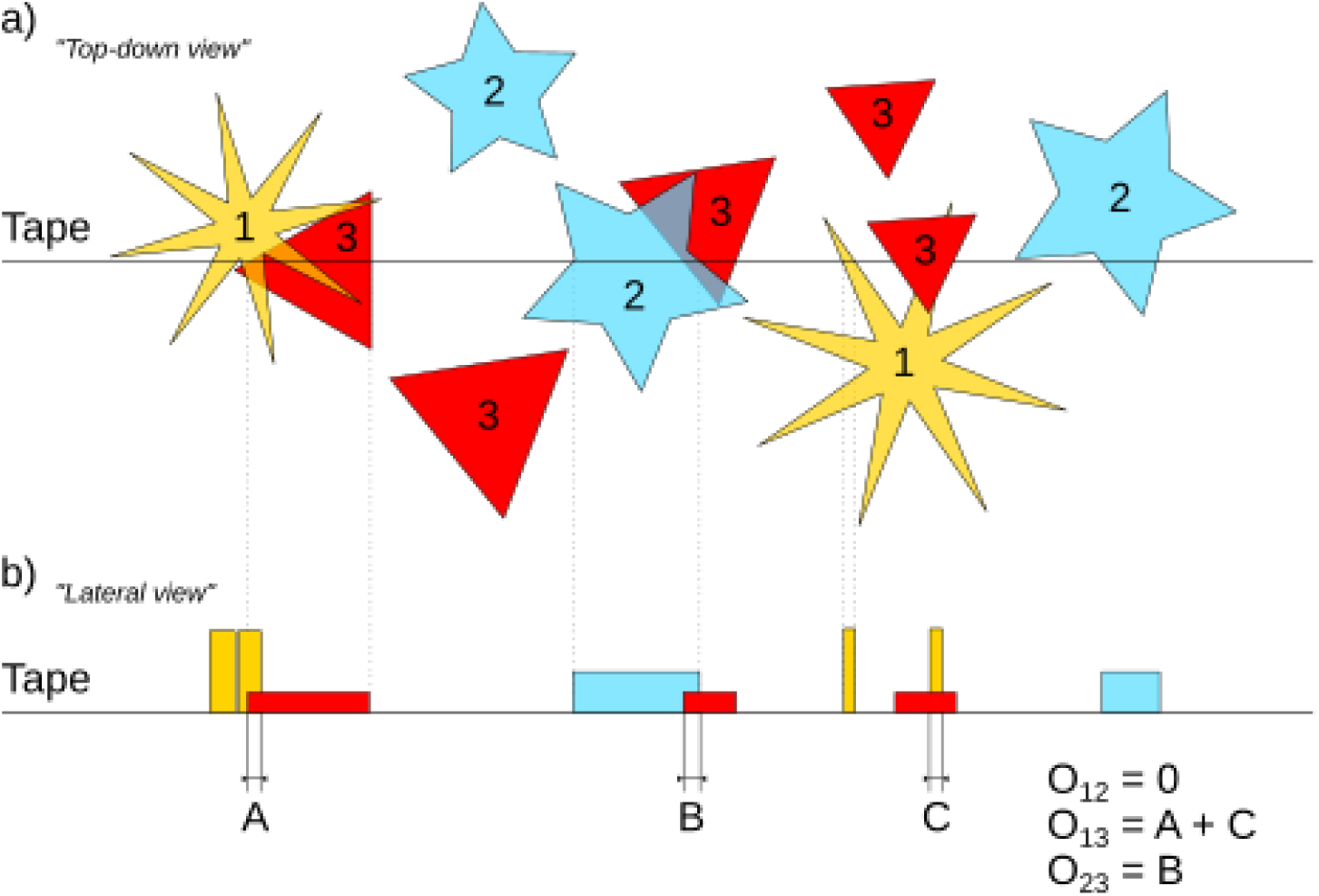
Measuring overlap between species using continuous transect data. In a), a top-down view of three plant species is given, with the straight line representing the line transect. For all individuals, the length over which they overlap is recorded (b), along with a class of height. The total overlap O_ij_ between all species is then computed, here O_{12} = 0 because these two species never overlap, O_13_ is equal to the sum of lengths A and C, as these species overlap twice, and O_23_ is equal to the length B. These overlap values are then compared to a null model where all species positions are randomized, i.e. rectangles in subfigure (b) are redistributed along the transect line to determine whether the pair of species is significantly associated.

**Table S1:** Trait names, descriptions and identifications (Trait ID) follow the thesaurus of plant characteristics, TOP (Garnier et al., 2017; accessible online at: https://www.top-thesaurus.org/), when available. Additional information were taken from Pérez-Harguindeguy et al. (2013)

| Trait | Abbreviation | Unit | Trait ID in TOP | Related function |
| --- | --- | --- | --- | --- |
| Leaf Dry Matter Content | LDMC | mg/g | TOP45 | Leaf tissue density, leaf physical resistance, stress tolerance |
| Specific Leaf Area | SLA | m<sup>2</sup>/kg | TOP50 | Photosynthetic rate, leaf longevity, relative growth rate |
| Leaf Area | LA | cm<sup>2</sup> | TOP25 | Light interception, energy balance |
| Leaf Carbon Content per unit mass | LCC<sub>m</sub> | mg/g | TOP452 | Conversion factor from gas exchange to mass based growth rate, lignin content |
| Leaf Nitrogen Content per unit mass | LNC<sub>m</sub> | mg/g | TOP462 | Light capture, photosynthetic rate |
| Leaf Phosphorus Content per unit mass | LPC<sub>m</sub> | mg/g | TOP463 | Energy acquisition, storage and utilization; component of nucleotides and nucleic acids |
| Leaf carbon isotopic fraction | Lδ<sup>13</sup>C | ‰ | Deviation of the leaf 13C/12C isotopic ratio | Gas exchange, water-use efficiency |
| Vegetative Plant Height | VPH | cm | TOP69 | Light capture, above-ground competition |
| Maximum horizontal plant length when plant in vegetative stage | D<sub>max</sub> | cm | the maximum horizontal length of aboveground parts of a plant measured for an undisturbed individual | Light capture, above-ground competition |
| Seed Mass | SeedMass | mg | TOP103 | Dispersal and regeneration strategy, seedling competition |

**Figure S4.**
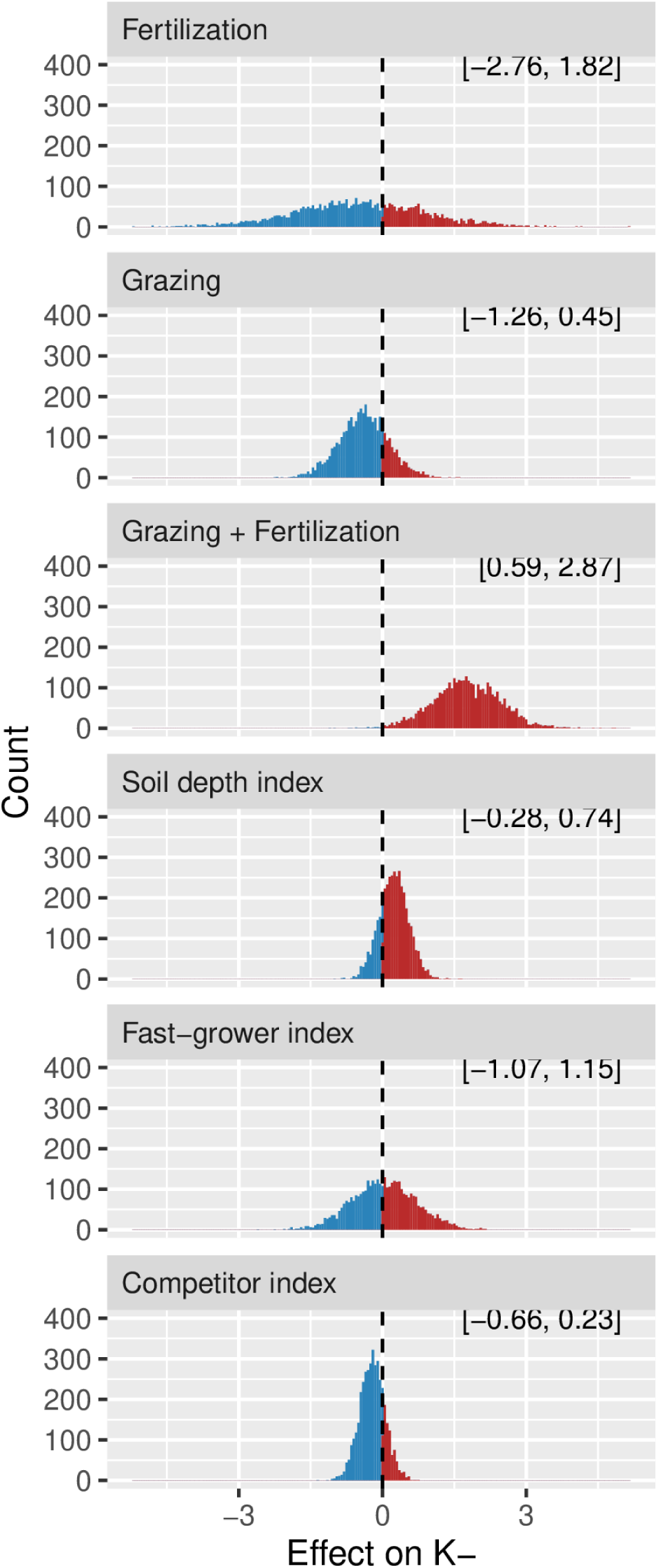
Posterior estimates for the linear model predicting K_SES_ as a function of Grazing, Fertilization, Soil depth index, and trait syndromes.

**Figure S5.**
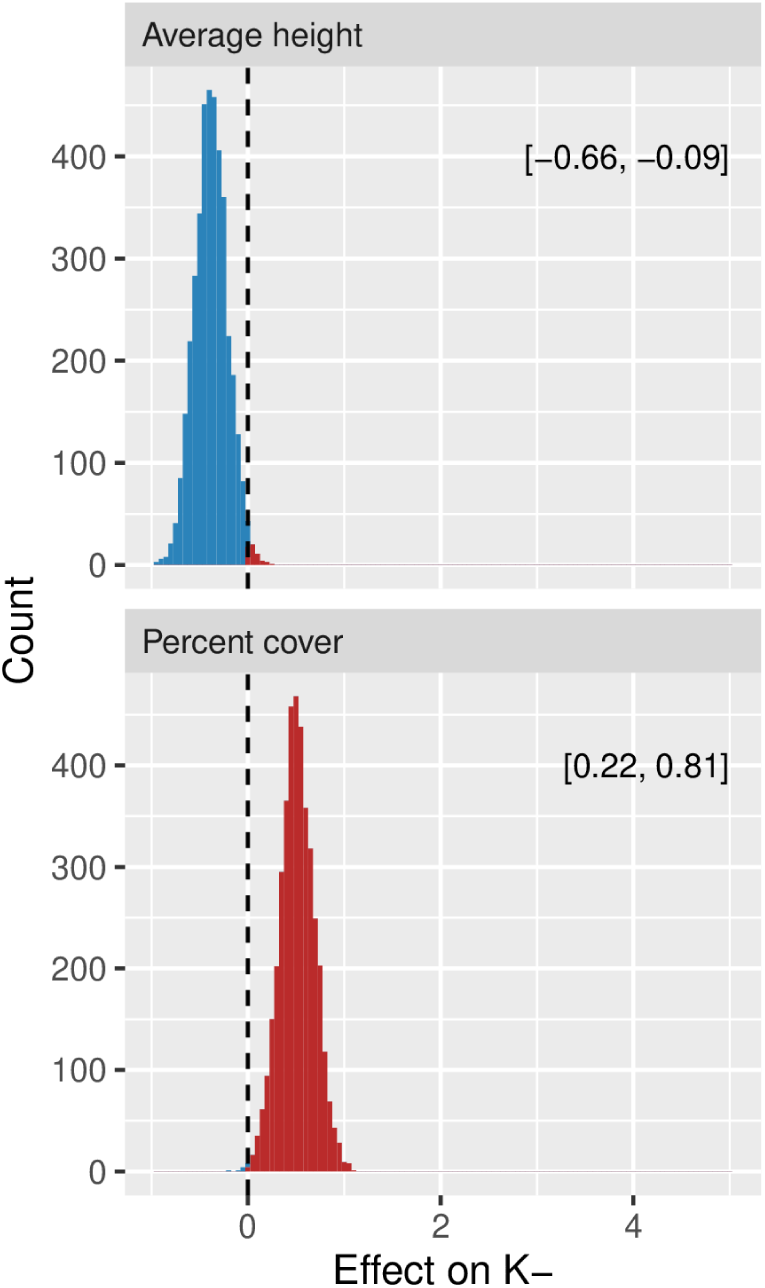
Posterior estimates for the linear model predicting K_SES_ as a function of average plant height and plant cover in a transect,

**Figure S6.**
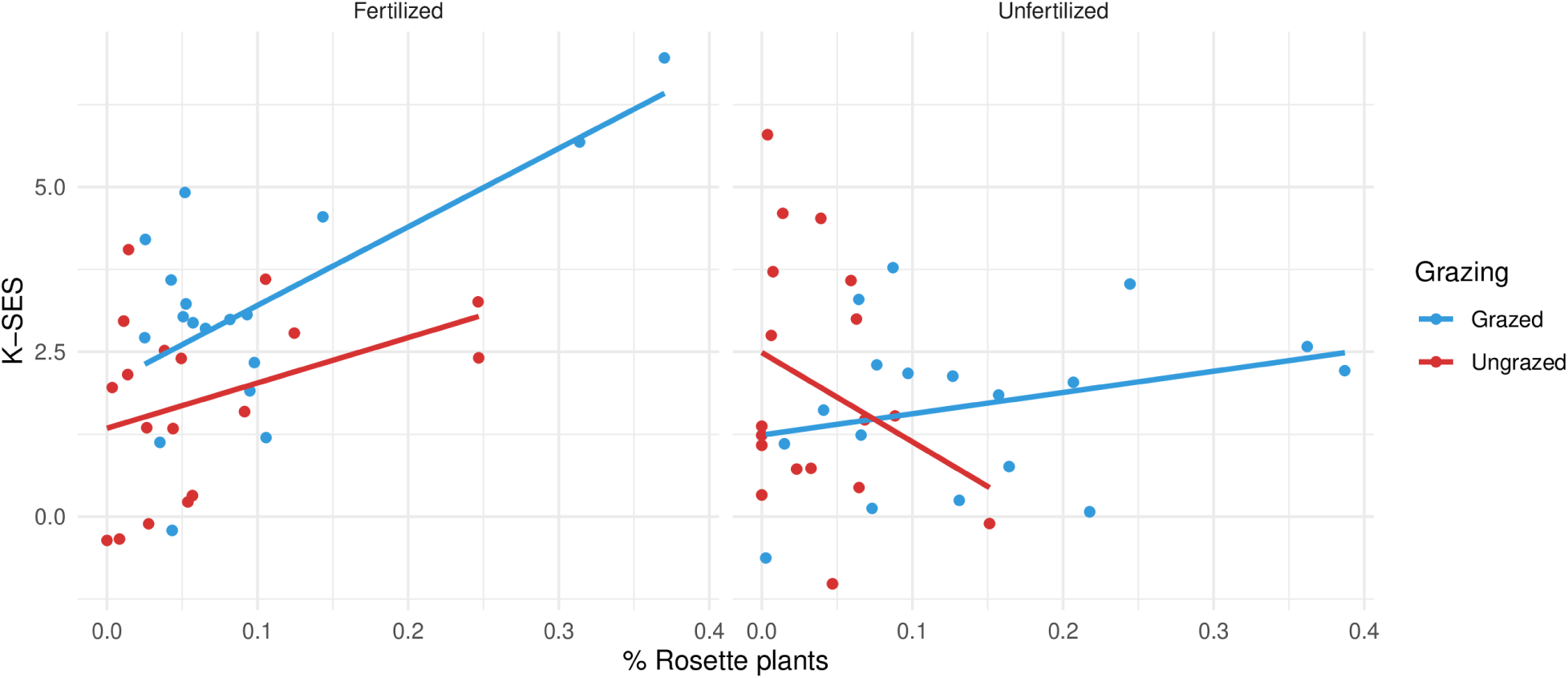
% Relationship between the % of plants with a rosette-like phenotype in the community, and K_SES_ for fertilized (panels) and grazing treatments (colors)

**Figure S7.**
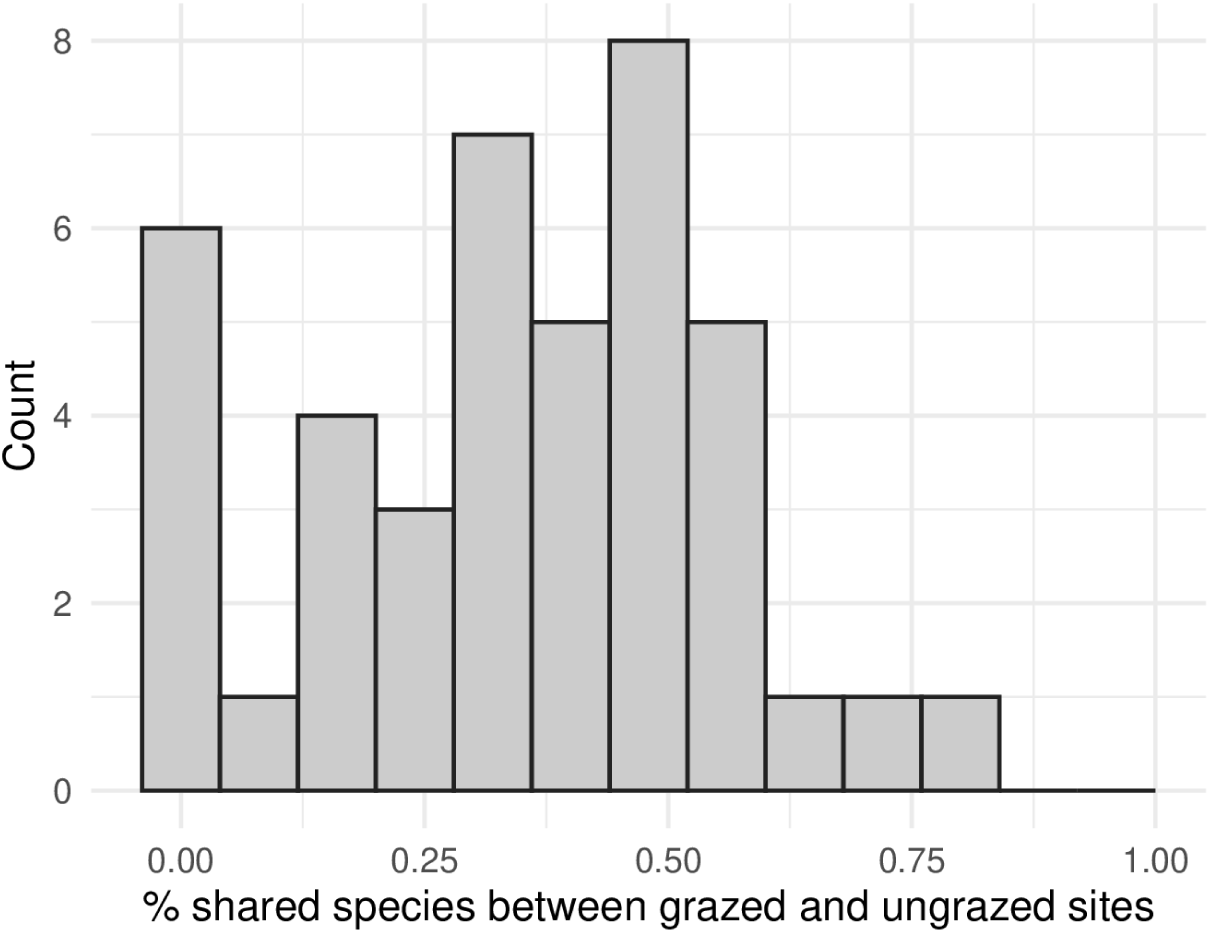
Histogram of the percentage of shared species between transects carried at the same location outside and inside exclosures

